# Decoding protein dynamicity in DNA ligase activity through deep learning-based structural ensembles

**DOI:** 10.1101/2024.11.07.622521

**Authors:** Nitesh Mishra, Sean Callaghan, Bryan Briney

## Abstract

Numerous proteins perform their functions by transitioning between various structures. Understanding the conformational ensembles associated with these states is essential for uncovering crucial mechanistic aspects that regulate protein function. In this study, we utilized AlphaFold3 (**AF3**) to investigate the structural dynamics and mechanisms of enzymes involved in DNA homeostasis, using NAD-dependent Taq ligases and human DNA Ligase 1 as a case example. Modifying the input parameters for AF3 yielded detailed conformational states of a DNA-binding enzyme, thereby offering enhanced mechanistic insights. We applied AF3 to model the various stages of thermophilic Taq DNA ligase activity, from its ground state to substrate-bound complexes, revealing significant mobility in the N-terminal adenylation and C-terminal BRCT domains. These detailed structural ensembles provided novel insights into the enzyme’s behavior during DNA repair, underscoring the potential of AF3 in elucidating mechanistic details critical for therapeutic and biotechnological targeting. Extending this approach to human LIG1, we examined its end-joining activity on double-strand breaks (**DSBs**) with short 3’ and 5’ overhangs. In alignment with published experimental data, AF3 conformational ensembles indicated LIG1 has lower catalytic efficiency for 5’ overhangs due to suboptimal DNA positioning within the catalytic site, demonstrating AF3’s capability to capture subtle yet functionally significant conformational differences by generating conformational ensembles capturing greater structural variance.

## INTRODUCTION

Proteins are indispensable biomolecules that perform a vast array of functions in living organisms. The advent of machine learning algorithms has greatly accelerated the pace of protein structure prediction. AlphaFold2 (**AF2**) brought a revolution to structural biology by accurately predicting single structures of proteins which ultimately led to prediction of nearly 200 million protein structures.^1–3^ However, the biological function of a protein often relies on multiple conformational substates that cannot be captured by a single static structural representation. Conformational changes, such as fold-switching and order-disorder transitions, are crucial for activity of numerous macromolecules and are prevalent throughout the proteome. Accurate modeling of protein dynamics within biological complexes is vital for understanding protein function and for the rational design of therapeutic agents. While AF2 does not predict conformational ensembles by default, several groups discovered that by subsampling the input multiple sequence alignments (**MSAs**), enabling dropouts, and increasing the number of predictions via random seed, it is possible to generate a series of structural predictions that capture various physiologically relevant conformations from the same sequence ^4–6^. Methods to predict conformational ensembles with AF2 have been established for small to moderate size proteins such as KaiB,^5^ AblI, and GM-CSF.^6^ Building on the success of AF2, the newest implementation of AlphaFold model, AlphaFold3 (**AF3**), now has the capability to jointly predict the structure of complexes that include proteins, nucleic acids, small molecules, ions, and modified residues, spanning the majority of the biomolecular space.^7^ AF3 demonstrates substantially improved accuracy for protein–nucleic acid interactions compared to many previous specialized tools with nucleic-acid-specific predictors, showcasing that high-accuracy modeling across biomolecular spaces is achievable within a single unified deep-learning framework.

Enzymes from hyperthermophiles, which live in high-temperature environments (over 80°C) and encode enzymes with unique thermophilic and thermostable properties, are a source of robust enzymes for biotechnological use that require DNA manipulation, such as molecular cloning, mutation detection, DNA assembly, and sequencing.^8–10^ One such enzyme, DNA ligase, is found in all three domains of life and is involved in DNA homeostasis to maintain DNA integrity via domain movements and active site remodeling. DNA ligases are critical in DNA repair, and are responsible for ligating DNA strands and aiding DNA replication, repair, and recombination within living cells. DNA ligases universally employ a three-step mechanism to catalyze phosphodiester bond formation,^11–14^ utilizing either ATP or NAD+ as cofactors. In the first step, a conserved lysine residue catalyzes nucleophilic attack on the 5′-phosphate of ATP or NAD+, forming a ligase-AMP intermediate and releasing pyrophosphate (**PPi**). In the second step, the AMP moiety is transferred to the 5′-phosphate of one DNA strand, generating an adenylated DNA intermediate. Finally, in the third step, a phosphodiester bond is formed between the 5′-phosphate of the adenylated DNA strand and the 3′-hydroxyl group of the second strand, releasing AMP as a byproduct. The binding of cofactors such as NAD or ATP initiates a transient closure in ligases to form the AMP-lysine phosphoramidate intermediate. The conserved domains within the core of DNA ligases are the adenylation domain (**AdD**) and the oligomer-binding domain (**OBD**). Certain ligases such as the hyperthermophilic bacterium *T. foliformi* contain additional domains including the BRCA1 C-terminus (**BRCT**) domain, a Zinc finger motif, and helix-hairpin-helix (**HhH**) motifs.^15^

To comprehensively investigate the potential of NAD-dependent ligases as robust molecular biological tools and as potential targets for antibacterial therapeutics, a thorough understanding of their structural dynamics and interactions during DNA repair is essential. DNA ligases recognize and repair damaged DNA ends via active site remodeling and given AF3’s ability to predict protein-nucleic acid interactions, we employed AF3 to investigate the conformational dynamics of thermophilic DNA ligase (Taq ligase) and human DNA ligase I (**LIG1**). By predicting the relative populations of Taq DNA ligase in its ground state, cofactor-bound state, and substrate + cofactor bound state in an ionic milieu, we aimed to identify major alternative conformations and their relative populations in a high-throughput manner.

## RESULTS

Our systematic modeling approach involved creating a comprehensive structural ensemble that captures various stages of Taq ligase activity, enhancing the statistical robustness of our findings and providing deeper insights into enzyme behavior. We generated 100 structural models (20 distinct seeds, 5 models per seed) for each of the five steps in DNA ligase-mediated repair of damaged DNA ends, for a total of 500 structures. We used different modeling parameters to simulate ground state (step 0), NAD-bound (step 1, representing cofactor binding), AMP-bound with metal ions (step 2, representing adenylate-intermediate), and DNA-bound states (steps 3a and 3b, capturing nicked and sealed DNA) (***Fig. 1***). This comprehensive modeling strategy allowed us to closely examine structural dynamics and protein/DNA interactions at each stage, elucidating the mechanism of Taq ligase-mediated DNA repair.

**Figure 1:**
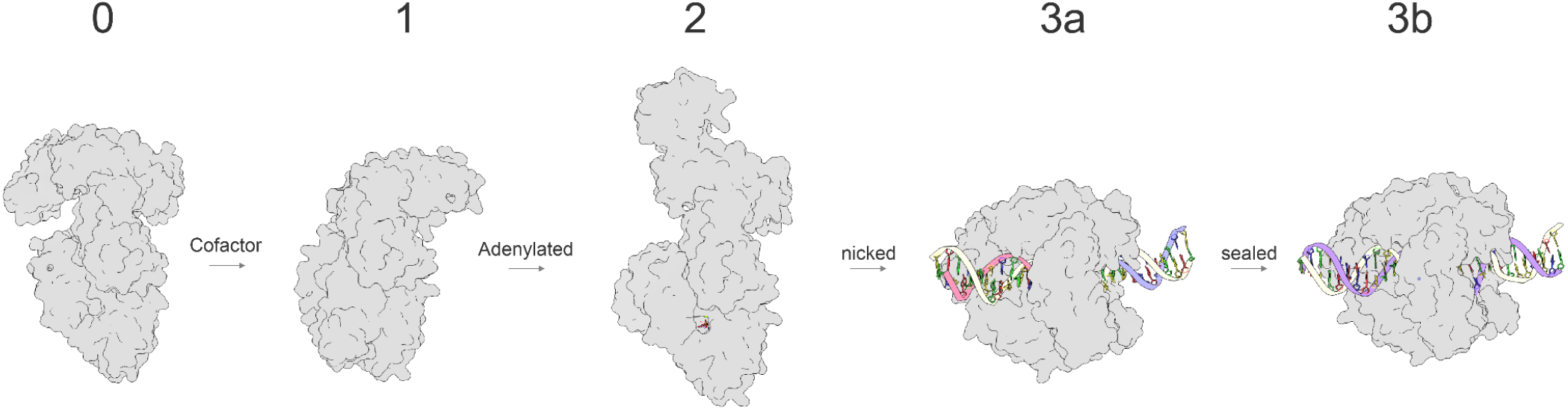
DNA Ligase Pathway. Schematic representation of all five stages that were modeled. Step 0 denotes the ground state of Taq DNA ligase, step 1 corresponds to pre-adenylated intermediate step, step 2 corresponds to post-adenylation step, step 3a and 3b correspond to nicked DNA at active site and ligated DNA stages.

### Step 0: Ground state

The conformational ensemble of Taq ligase in its ground state (*step 0*) showed significant mobility at the hinges for the N-terminal adenylation and C-terminal BRCT domains (***Fig. 2A and S1A***), two regions crucial for cofactor binding and DNA regulation, respectively. The core protein subdomains were predicted with high confidence across all structures, with the adenylation domain and the BRCT domain also showing good confidence despite the dynamicity at the hinges that connect all subdomains (***Fig. S1A***). In *T. filiformi* DNA ligase, the BRCT domain is known to play a role in regulating DNA binding and release. The conformational ensemble of this domain exhibits considerable mobility in the open conformation (***Fig. S1B***), as expected for its function as a gate to control DNA binding and release. Confidence scores (pLDDT scores) at the hinges connecting the AdD and BRCT domains to the core were relatively lower and can be attributed to the structurally dynamic role of the two domains, as evident by high confidence scores for hinges connecting BRCT domain in DNA bound form where its mobility is significantly reduced (***Fig. S1C***). The core domains are typically more rigid to ensure consistent interactions with DNA and other molecules, thereby minimizing conformational variability. In line with this observation, the core domain containing the OBD and zinc binding site showed the lowest degree of fluctuations in the ensemble, suggestive of their role in maintaining structural integrity (***Fig. 2A and S1D***).

**Figure 2.**
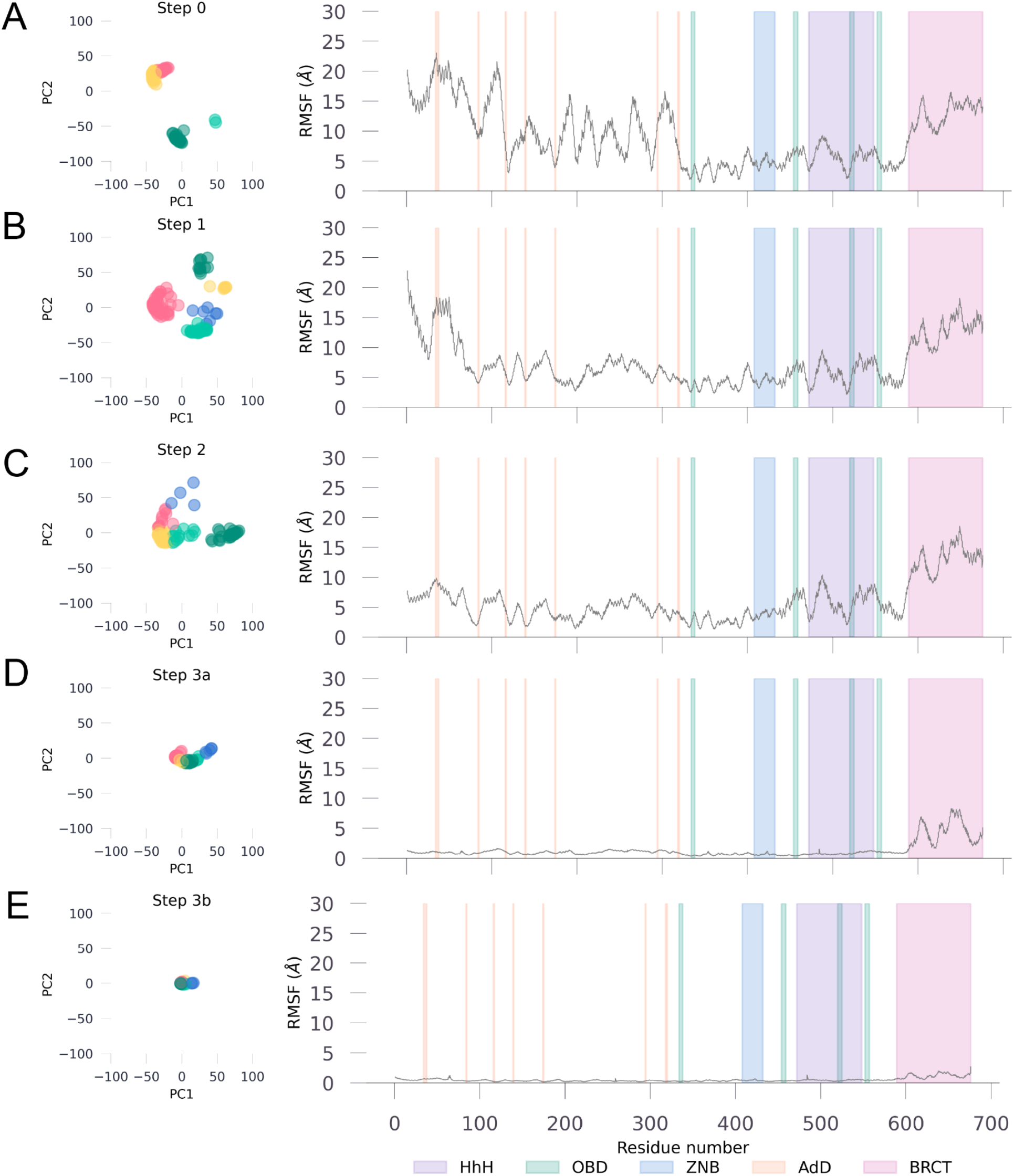
Domain-Specific Backbone PCA and RMS Fluctuations in Taq Ligase. Left panel shows the Taq ligase backbone at that step reduced to 2 principal components for all models within the ensemble. Right panel shows the backbone fluctuations at each residue (root-mean square fluctuations) compared to an average structure calculated using all models from the ensemble at that step. Domains are color coded per the key given at the bottom right corner.

### Step 1: Cofactor-bound state

Our structural modeling of Taq ligase with NAD (step 1) predominantly resulted in models adopting a closed conformation, consistent with reported transient closures in similar ligases.^15^ In this step, ATP is used to form a high-energy lysine-AMP intermediate, and all models in the ensemble had the AMP moiety next to the catalytic lysine (K118) (***Fig. 2B and 3A-B***). AF3 currently does not allow the adenylate modifications of lysine, but the placement of AMP moiety is in alignment with several structures of ATP-dependent ligases as well as *E. coli* ligase (**LigA**).^10,16^

### Step 2: Adenylate-intermediate state

Modeling Taq ligase in complex with AMP and metal ions (step 2), predominantly showed open conformations (***Fig. 3C***), facilitating an accessible binding surface for damaged DNA after enzymatic activity.

**Figure 3.**
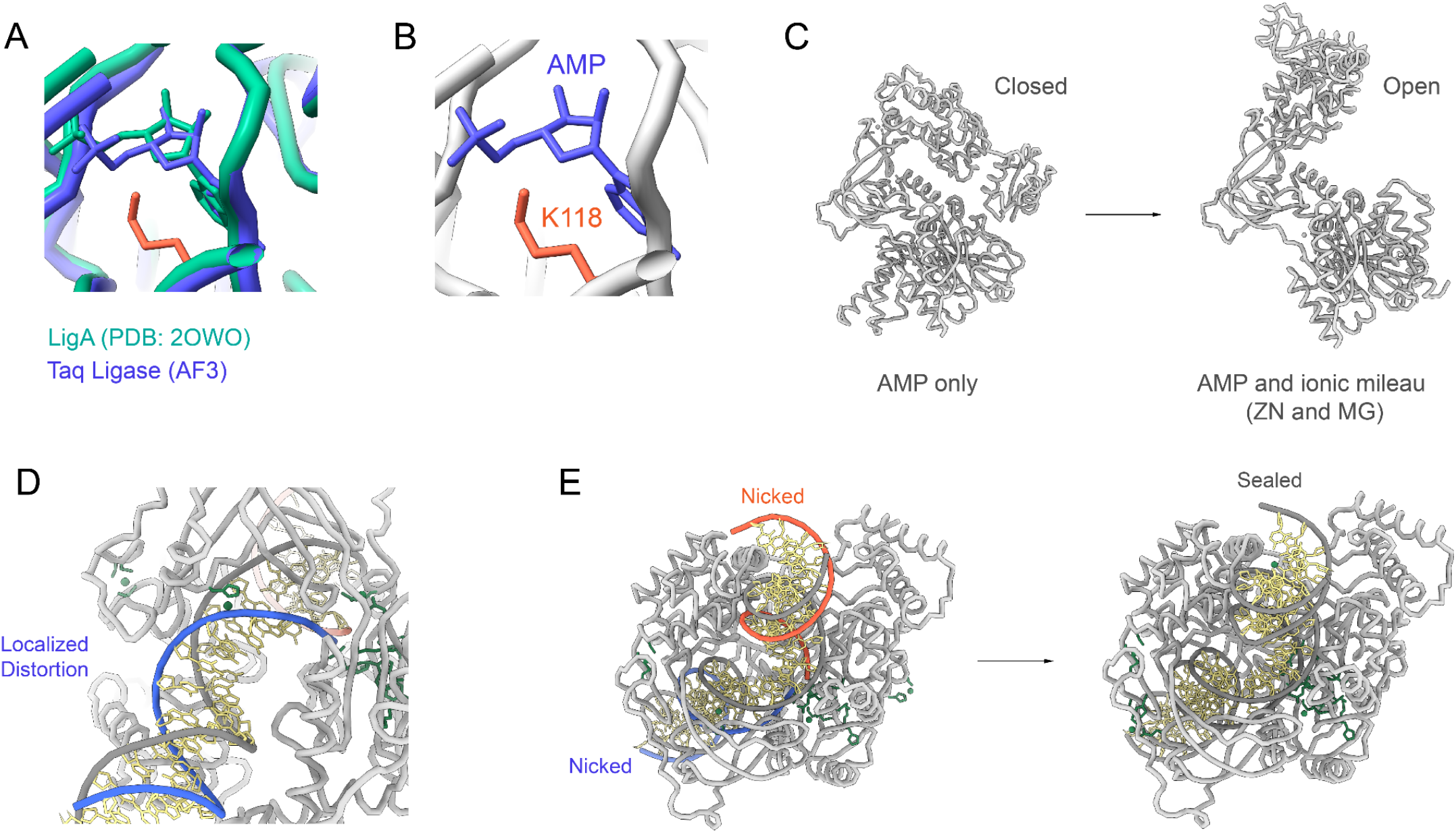
Conformational Changes and DNA Interactions in Taq Ligase Structural Ensemble. (A) Closed conformation of Taq ligase with NAD, showing AMP moiety adjacent to catalytic lysine (K118) in the ensemble. (B) Zoomed view of the catalytic site in the closed state, highlighting AMP’s position next to K118. (C) Closed-to-Open conformation transition of Taq ligase in the adenylate-intermediate state, allowing an accessible surface for DNA binding. (D) Closed conformation with nicked DNA, depicting the formation of a ternary complex for efficient phosphodiester bond catalysis. (E) DNA adopts an RNA-like A-form helix near the nick in the LigA complex, reflecting localized structural distortions during repair.

### Step 3a and 3b: Capturing nicked and sealed DNA

Structural studies of DNA ligases, such as the ChVLig-AMP complex, reveal significant domain rearrangements upon DNA binding, highlighting key structures and interactions essential for therapeutic targeting and enzyme design in biotechnology.

Comparisons of ligase complexes across bacterial species underscore massive domain shifts, ranging from 50 to 90 Å, during substrate binding and catalysis. Each ligation step involves distinct domains, with the NTase domain universally required across all stages. The structural transition observed during DNA binding and clamp formation by LigA involved a significant rotation of the OB domain, optimizing interaction with DNA’s minor groove in a near 180° rotation. In step 3a, modeling the complex with nicked DNA showed closed conformations conducive to forming a ternary complex essential for efficient phosphodiester bond catalysis. Interactions between LigA and DNA near the nick showed a localized distortion (***Fig. 3D-E***), causing the DNA to adopt an RNA-like A-form helix consistently across the structural ensemble. This RNA-like conformational change highlights structural rearrangements within the DNA during the nick repair process, suggesting a pivotal role in maintaining DNA integrity through specific interactions. These findings underscore the intricate mechanisms by which NAD-dependent ligases, like Taq ligase, contribute to DNA repair and highlight their potential as precise targets for developing antibacterial therapies, mitigating risks to human DNA repair mechanisms.

### Modeling double-strand break repair by LIG1

Building on our observations of the predicted conformational dynamics of Taq ligase, we extended our simulations to human DNA Ligase I (LIG1) to further validate the robustness and applicability of this approach across different ligase types. This extension aimed to explore whether AF3 could similarly capture the structural nuances and mechanistic details of LIG1, particularly in contexts where its catalytic efficiency varies.^17,18^ We focused on the activity of LIG1 in repairing double-strand breaks (DSBs) with different overhangs as a means of investigating how the conformational adaptations of LIG1 influence enzymatic efficiency. This comparative approach not only tested the versatility of AF3 in modeling diverse ligases but also provided deeper insights into the specific challenges faced by LIG1 in mediating DNA end-joining, thereby broadening our understanding of DNA repair mechanisms and informing potential therapeutic strategies.

The ligation activities of LIG1 for the repair of single-strand breaks (SSBs) have been previously characterized, highlighting its crucial role in maintaining genome integrity.^11,19–21^ For the end-joining of DSBs that lack cohesive ends, DNA ligases must overcome the additional challenge of properly positioning two DNA molecules to catalyze the formation of the phosphodiester bond without the aid of intermolecular base pairing interactions.^18^

Recent findings demonstrated that LIG1 can perform end-joining of DNA ends but exhibits lower catalytic efficiency when the substrate has short 5’ overhangs.^18^

To further investigate this phenomenon, we simulated LIG1-mediated ligation of a 30mer DNA with short 3’ overhangs (3DSB) or short 5’ overhangs (5DSB) (***Fig. 4A***) by generating conformational ensembles using AF3. These ensembles revealed notable differences in the positioning and interaction of the DNA substrates with LIG1.

**Figure 4.**
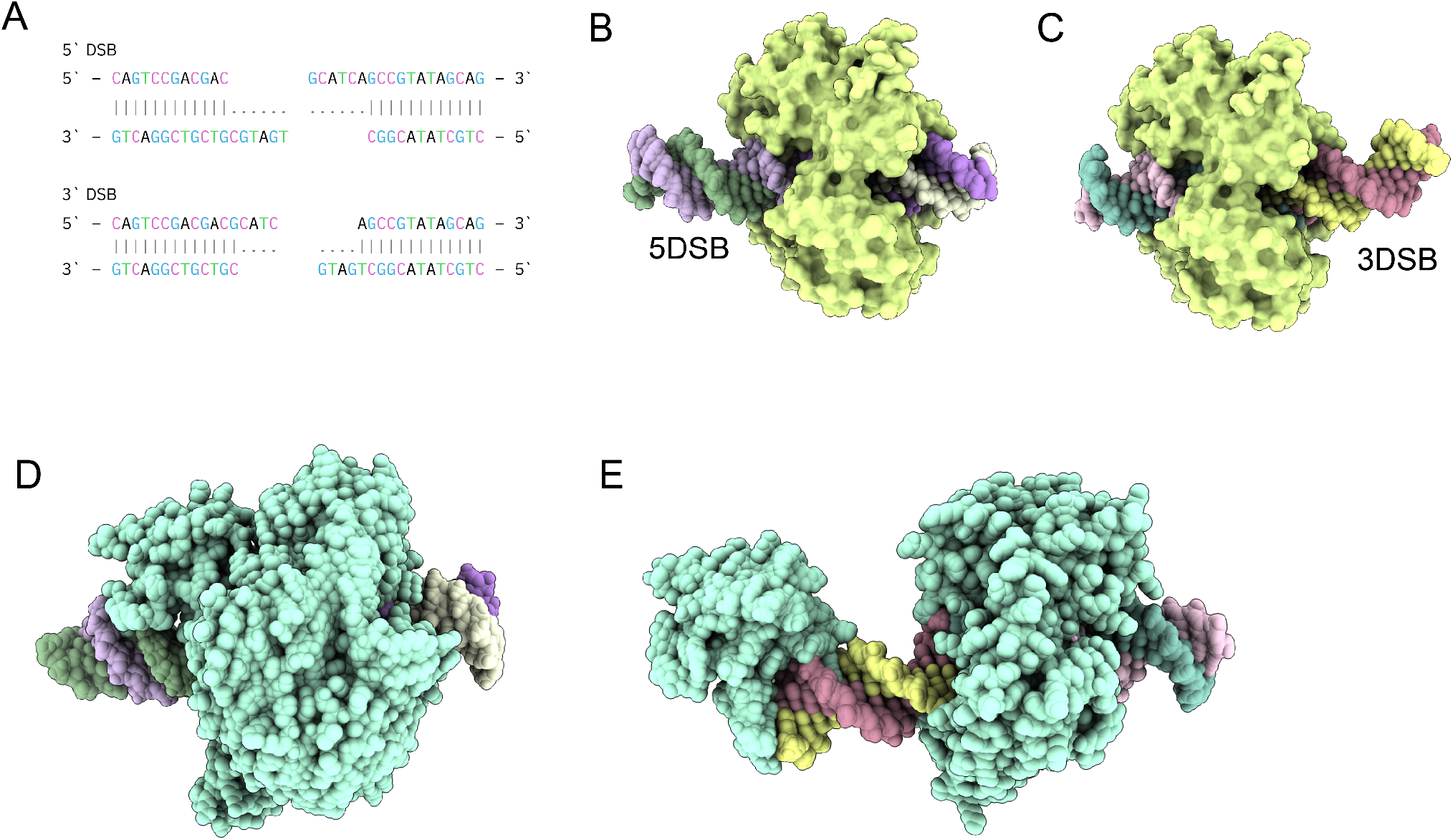
Conformational Dynamics of Human DNA Ligase I (LIG1) in Double-Strand Break Repair. (A) 30mer DNA sequences with short 3’ (3DSB) and 5’ overhangs (5DSB) used for modeling LIG1 mediated ligation. (B) Conformational ensemble of LIG1 with 5DSB, illustrating suboptimal DNA positioning with a protruding end and reduced contact area with the enzymatic site. (C) Conformational ensemble of LIG1 with 3DSB, demonstrating optimal DNA positioning centrally within the catalytic site, facilitating efficient catalysis. (D) Structural representation of the LIG3b-5DSB complex, showing the DNA enveloped in the central cavity of LIG3b. (E) Structural representation of the LIG3b-3DSB complex, with a large protruding end stabilized by the N-terminal zinc finger domain of LIG3b.

In the conformational ensemble for LIG1 with 5DSB, the DNA was positioned suboptimally, with a larger protruding end on one side and less contact area with the enzymatic site of LIG1 (***Fig. 4B***). In contrast, for 3DSB, the DNA was observed to be centrally located within the catalytic site (***Fig. 4C***), presumably facilitating efficient catalysis. These observations provide structural insights into the mechanism underlying lower catalytic activity of LIG1 for substrates with 5’ overhangs. The reduced contact area and suboptimal positioning of the DNA within the catalytic site likely hinder the enzyme’s ability to effectively catalyze the ligation reaction, providing a structural rationale for LIG1’s lower catalytic efficiency with 5DSB. This conformational difference underscores the importance of substrate positioning and enzyme-DNA interactions in determining the efficiency of the ligation process. We next simulated the 5DSB and 3DSB with LIG3b, which has been shown to have similar catalytic activity for both types of overhangs.^18^ In contrast with LIG1, 5DSB was enveloped in the central cavity within the LIG3b-5DSB complex (***Fig. 4D***) while LIG3b-3DSB had a large protruding end that was stabilized by the N-terminal zinc finger of LIG3b (***Fig. 4E***). The N-terminal zinc finger (ZnF) domain of LIG3b is homologous to those of poly (ADP-ribose) polymerase 1 (PARP-1). This N-terminal region of LIG3 appears to be dynamic and has been proposed to serve as a nick sensor for SSB ligation, as well as to play an essential role in LIG3-catalyzed DNA end joining.

## DISCUSSION

In this study, we sought to harness the advanced capabilities of AF3 to elucidate the structural dynamics of DNA ligase, a crucial enzyme in DNA replication, repair, and recombination across all domains of life. Recognizing the importance of conformational ensembles for the biological function of proteins, our research aimed to extend the predictive power of AF3 beyond single structures to capture multiple conformational states representing the entire catalytic activity cycle of DNA ligase.

The application of deep learning methods, exemplified by AF3, has profoundly enhanced our ability to explore the intricate structural mechanisms underlying enzymatic processes such as DNA repair mediated by NAD-dependent ligases. By generating detailed structural ensembles at various stages of Taq ligase activity, we gained significant insights into the dynamic interactions and conformational changes crucial for catalysis. AF3’s predictive accuracy has not only provided a comprehensive view of ligase behavior from ground state to substrate-bound complexes but has also illuminated novel structural features and interactions that were previously inaccessible. These insights pave the way for targeted drug design aimed at disrupting bacterial DNA repair mechanisms while sparing human counterparts, thereby offering new avenues for combating antibiotic resistance and advancing precision medicine. In essence, the integration of deep learning approaches in structural biology not only deepens our fundamental understanding of enzyme function but also holds promise for transformative innovations in therapeutic development and biotechnological applications

By leveraging AF3’s ability to model protein-nucleic acid interactions with high accuracy, we generated structural ensembles of Taq ligase across various stages of the DNA repair process. Our approach involved detailed modeling of each step in the ligase-mediated repair of damaged DNA ends, providing comprehensive insights into the enzyme’s conformational changes and interactions. This study not only advances our understanding of DNA ligase’s mechanism but also demonstrates the potential of next-generation AI tools like AF3 to predict complex biomolecular interactions, paving the way for future applications in biotechnology and therapeutic development. We reveal that AF3 successfully predicts different conformations of thermophilic Taq ligase in its ground state, transition state, and active state. This underscores the potential of AF3 in elucidating the dynamic nature of protein conformations, providing deeper insights into their mechanistic roles in various biological processes. The framework outlined in this study is designed to be generalizable and can be applied to gain mechanistic insights into various unique enzymes, such as the Sulfophobococcus zilligii DNA ligase, which can utilize multiple substrates including ATP, ADP, and GTP. The structural insights into the ligase mechanism and substrate specificity garnered from our research offer promising avenues for future studies on enzyme evolution.

The implications of these findings extend beyond the basic understanding of LIG1’s catalytic mechanism. They suggest that enhancing the interaction between LIG1 and DNA substrates with 5’ overhangs could be a potential strategy to improve the efficiency of end-joining repair. This could be achieved through protein engineering or the development of small molecules that stabilize the enzyme-substrate complex. Additionally, these insights could inform the design of more effective DNA repair enzymes for therapeutic applications, particularly in the context of genetic diseases or cancer where DSB repair is critical. Furthermore, the ability of AF3 to generate detailed conformational ensembles that capture these subtle yet significant differences highlight its potential as a powerful tool in structural biology. By providing a deeper understanding of enzyme-substrate interactions, AF3 can aid in the rational design of enzymes with enhanced catalytic properties, contributing to advances in biotechnology and medicine. Overall, these ensembles not only elucidate the specific challenges faced by LIG1 in end-joining DSBs with 5’ overhangs but also demonstrate the broader utility of deep learning methods in uncovering the intricacies of enzymatic mechanisms.

Once the weights for AlphaFold3 are released, by adjusting the input parameters, researchers can exploit AF3’s capabilities to explore the conformational landscape of enzymes more comprehensively by modifying the input ligands and cofactors. This approach facilitates the identification of transient intermediates and conformational shifts that are often missed by traditional methods. The ability to predict such detailed states not only deepens our understanding of the enzyme’s mechanism but also aids in the identification of potential targets for therapeutic intervention. By leveraging these insights, it may be possible to design novel catalysts that enhance our understanding of the genetics and cell biology of DNA repair. Moreover, these advancements could significantly impact the practice of enzymatic joining and the synthesis of nucleic acids, leading to innovative applications in biotechnology and medicine. The current study not only provides a deeper understanding of NAD-dependent ligases but also sets the stage for future research aimed at exploiting these findings for the development of new biotechnological tools and therapeutic strategies. Additionally, these ensembles have the potential to serve as a rich dataset for rational protein engineering to make robust DNA ligases.

## METHODS

### Overview of the structural prediction pipeline

For step 1, we used only the amino acid sequence of Taq ligase (UniProt: P26996), representing the enzyme in its ground state. In step 2, the model included the amino acid sequence along with NAD as a ligand. Step 3 involved the amino acid sequence with AMP as a ligand, along with Zn and Mg ions. For step 4, we utilized the template (sense) DNA strand and the non-template (anti-sense) strands as two separate inputs. Steps 2 to 4 represent the transition states of the enzyme during the repair process. Finally, in step 5, the two nicked DNA strands were provided as a continuous sequence, simulating the final DNA ligation process. All structures were predicted using the AlphaFold3 server with 24 different seeds (5 model output per seed, 120 structural models for each step). This comprehensive modeling approach allowed us to closely examine the structural dynamics and interactions at each stage of the DNA repair process, providing detailed insights into the mechanism of Taq ligase-mediated DNA end repair.

### Structural analysis

PCA and RMSF calculations were carried out in python using the MDAnalysis^22,23^ package.

### LIG-1 ensemble

Structure model for LIG1 in AlphaFold database has the first 261 residues as disordered, therefore, we used residue 261 - 919 for generating the conformational ensemble. For 3DSB, we used the following nucleotide sequence (sense strand: CAGTCCGACGACGCATC (break) AGCCGTATAGCAG; anti-sense strand: CTGCTATACGGCTGATG (break) CGTCGTCGGACTG). For 5DSB, we used the following nucleotide sequence (sense strand: CAGTCCGACGAC (break) GCATCAGCCGTATAGCAG; anti-sense strand: CTGCTATACGGC (break) TGATGCGTCGTCGGACTG). Ensembles were generated with MG ions and AMP as ligands (using 24 different seeds).

### Protein Structure Visualization

For visualizing aligned structures, generating superimposed ensemble, color structures by AlphaFold3 confidence score (**pLDDT**), UCSF ChimeraX^24^ (version 1.8) was used.

## Data and Code Availability

All json inputs files used and the output raw data from AlphaFold server, and code used for PCA and RMSF calculations in this study are available on Zenodo (doi: 10.5281/zenodo.14048516) and on GitHub (github.com/brineylab/taq-ligase-structural-ensemble).

## AUTHOR CONTRIBUTIONS

Conceptualization: NM

Code and modeling: NM, SC

Data analysis: NM, SC

Manuscript preparation and revisions: NM, SC, BB

---

**Figure S1:**
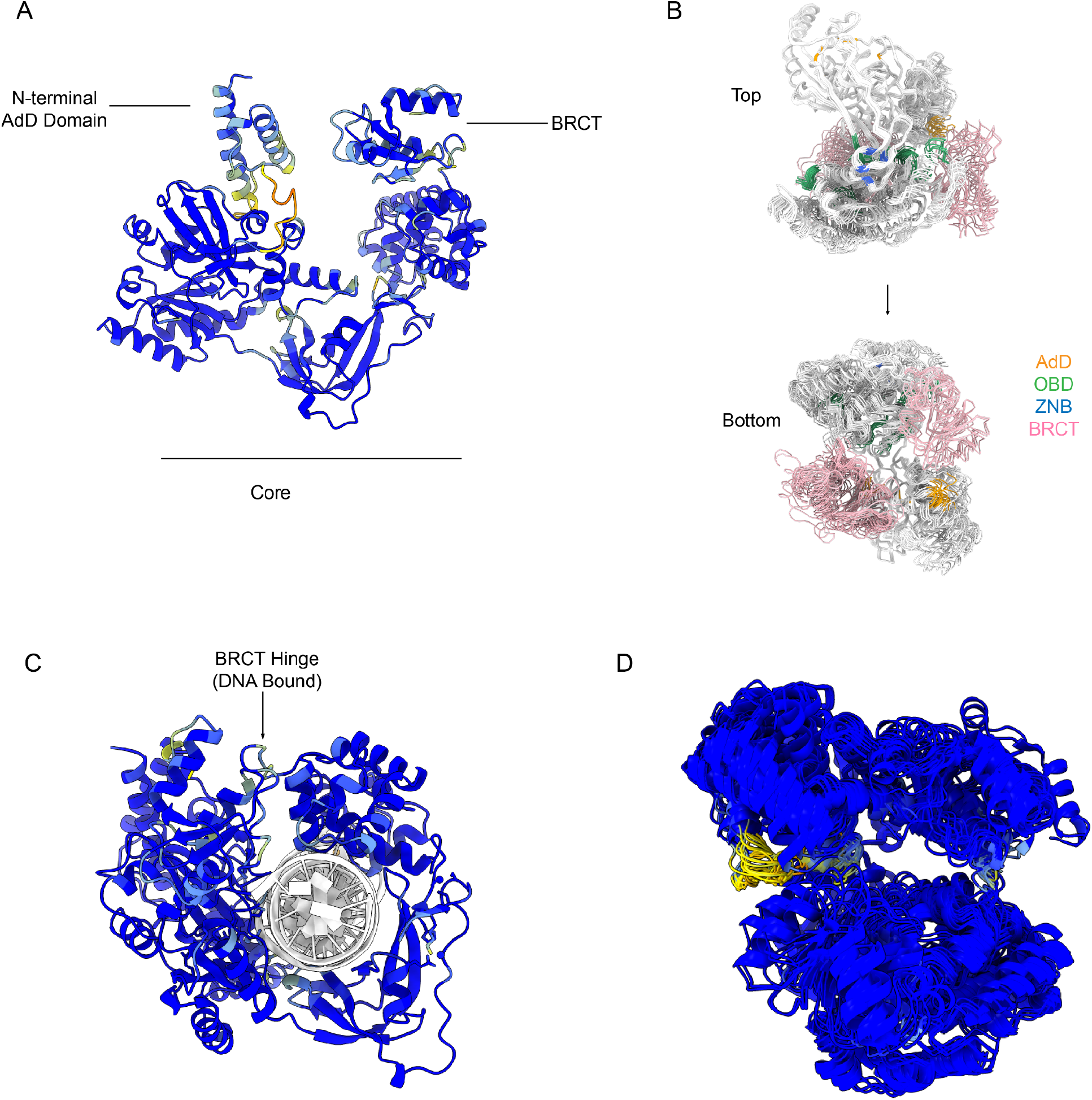
Domain-Specific Confidence and Structural Flexibility in Taq Ligase. (A) Confidence score across domains. (B) Conformational flexibility highlighted by superimposing the ensemble. (C) Hinge confidence in DNA bound form. (D) High confidence in the core domain across the ensemble.

## REFERENCES

1. Jumper, J. et al. Highly accurate protein structure prediction with AlphaFold. Nature 596, 583–589 (2021).

2. Tunyasuvunakool, K. et al. Highly accurate protein structure prediction for the human proteome. Nature 596, 590–596 (2021).

3. Varadi, M. et al. AlphaFold Protein Structure Database in 2024: providing structure coverage for over 214 million protein sequences. Nucleic Acids Res. 52, D368–D375 (2024).

4. Del Alamo, D., Sala, D., Mchaourab, H. S. & Meiler, J. Sampling alternative conformational states of transporters and receptors with AlphaFold2. eLife 11, e75751 (2022).

5. Wayment-Steele, H. K. et al. Predicting multiple conformations via sequence clustering and AlphaFold2. Nature 625, 832–839 (2024).

6. Monteiro da Silva, G., Cui, J. Y., Dalgarno, D. C., Lisi, G. P. & Rubenstein, B. M. High-throughput prediction of protein conformational distributions with subsampled AlphaFold2. Nat. Commun. 15, 2464 (2024).

7. Abramson, J. et al. Accurate structure prediction of biomolecular interactions with AlphaFold 3. Nature 630, 493–500 (2024).

8. Verma, D., Kumar, V. & Satyanarayana, T. Genomic attributes of thermophilic and hyperthermophilic bacteria and archaea. World J. Microbiol. Biotechnol. 38, 135 (2022).

9. Vieille, C. & Zeikus, G. J. Hyperthermophilic enzymes: sources, uses, and molecular mechanisms for thermostability. Microbiol. Mol. Biol. Rev. MMBR 65, 1–43 (2001).

10. Shi, J., Oger, P. M., Cao, P. & Zhang, L. Thermostable DNA ligases from hyperthermophiles in biotechnology. Front. Microbiol. 14, 1198784 (2023).

11. Pascal, J. M. DNA and RNA ligases: structural variations and shared mechanisms. Curr. Opin. Struct. Biol. 18, 96–105 (2008).

12. Shuman, S. DNA ligases: progress and prospects. J. Biol. Chem. 284, 17365–17369 (2009).

13. Williamson, A., Grgic, M. & Leiros, H.-K. S. DNA binding with a minimal scaffold: structure-function analysis of Lig E DNA ligases. Nucleic Acids Res. 46, 8616–8629 (2018).

14. Williamson, A. & Leiros, H.-K. S. Structural insight into DNA joining: from conserved mechanisms to diverse scaffolds. Nucleic Acids Res. 48, 8225–8242 (2020).

15. Lee, J. Y. et al. Crystal structure of NAD(+)-dependent DNA ligase: modular architecture and functional implications. EMBO J. 19, 1119–1129 (2000).

16. Shi, K. et al. T4 DNA ligase structure reveals a prototypical ATP-dependent ligase with a unique mode of sliding clamp interaction. Nucleic Acids Res. 46, 10474–10488 (2018).

17. McNally, J. R. & O’Brien, P. J. Kinetic analyses of single-stranded break repair by human DNA ligase III isoforms reveal biochemical differences from DNA ligase I. J. Biol. Chem. 292, 15870–15879 (2017).

18. McNally, J. R., Ames, A. M., Admiraal, S. J. & O’Brien, P. J. Human DNA ligases I and III have stand-alone end-joining capability, but differ in ligation efficiency and specificity. Nucleic Acids Res. 51, 796–805 (2023).

19. Taylor, M. R., Conrad, J. A., Wahl, D. & O’Brien, P. J. Kinetic mechanism of human DNA ligase I reveals magnesium-dependent changes in the rate-limiting step that compromise ligation efficiency. J. Biol. Chem. 286, 23054–23062 (2011).

20. Liddiard, K. et al. DNA Ligase 1 is an essential mediator of sister chromatid telomere fusions in G2 cell cycle phase. Nucleic Acids Res. 47, 2402–2424 (2019).

21. Kamble, P., Hall, K., Chandak, M., Tang, Q. & Çağlayan, M. DNA ligase I fidelity mediates the mutagenic ligation of pol β oxidized and mismatch nucleotide insertion products in base excision repair. J. Biol. Chem. 296, 100427 (2021).

22. Michaud-Agrawal, N., Denning, E. J., Woolf, T. B. & Beckstein, O. MDAnalysis: a toolkit for the analysis of molecular dynamics simulations. J. Comput. Chem. 32, 2319–2327 (2011).

23. Gowers, R. et al. MDAnalysis: A Python Package for the Rapid Analysis of Molecular Dynamics Simulations. in 98–105 (Austin, Texas, 2016). doi:10.25080/Majora-629e541a-00e.

24. Meng, E. C. et al. UCSF ChimeraX: Tools for structure building and analysis. Protein Sci. Publ. Protein Soc. 32, e4792 (2023).

